# Multiplexed optical barcoding and sequencing for spatial omics

**DOI:** 10.1101/2024.06.04.597495

**Authors:** Aditya Venkatramani, Didar Ciftci, Limor Cohen, Christopher Li, Khanh Pham, Xiaowei Zhuang

**Affiliations:** Howard Hughes Medical Institute, Department of Chemistry and Chemical Biology, Department of Physics, Harvard University, Cambridge, MA 02138, USA

## Abstract

Spatial omics has brought a fundamental change in the way that we study cell and tissue biology in health and disease. Among various spatial omics methods, genome-scale imaging allows transcriptomic, 3D-genomic, and epigenomic profiling of individual cells with high spatial (subcellular) resolution but often requires a preselection of targeted genes or genomic loci. On the other hand, spatially dependent barcoding of molecules followed by sequencing provides untargeted, genome-wide profiling but typically lacks single-cell resolution. Here, we report a spatial omics method that could potentially combine the power of the two approaches by optically controlled spatial barcoding followed by sequencing. Specifically, we utilize patterned light to encode the locations of cells in tissues using oligonucleotide-based barcodes and then identify the barcoded molecular content, such as mRNAs, by sequencing. This optical barcoding method is designed with multiplexing and error-correction capacity and achieved by a light-directed ligation chemistry that attaches distinct nucleic-acid sequences to the reverse transcribed cDNA products at different locations. As a proof of principle for this method, we demonstrated high-efficiency in situ light-directed ligation, spatially dependent barcoding with multiplexed light-controlled ligations, and high-accuracy detection of spatially barcoded mRNAs in cells.

## Introduction

Spatial omics provides a systematic approach to measuring expression profiles and molecular signatures of cells, identifying cell types and states in their native environment, and mapping the spatial organizations and interactions of distinct cell types in tissues. Recent advances in spatial omics have enabled the measurements of the transcriptome, 3D-genome, epigenome, and protein expression profiles of cells in tissues [1-3]. These measurements have generated detailed cell-type atlases of a variety of organs and whole organisms at various stages of development, revealed changes in gene expression patterns in disease states, and provided comprehensive resources to study molecular interactions amongst different cell types [1, 2, 4]. The rapid pace of knowledge and hypothesis generation by this approach has, in turn, provided an incentive to develop and improve spatial omics methods constantly.

Spatial omics methods fall broadly into two categories: approaches based on massively multiplexed imaging and approaches based on next-generation sequencing. Imaging-based methods allow simultaneous detection, quantification, and localization of thousands of RNA species and/or genomic loci, as well as dozens of proteins, with single-cell and subcellular resolution [2, 3]. However, most imaging-based methods require the pre-selection of genes and genomic loci. On the other hand, sequencing-based methods do not require a pre-selection of targeted molecules and provide genome-wide information, but with lower spatial resolution than imaging-based methods. These sequencing-based methods generally rely on spatially dependent barcoding of molecules in samples using distinct DNA sequences, followed by extraction and sequencing of the molecular content to determine both the genetic identity and spatial locations of molecules [1, 3]. Because such spatial barcoding occurs in two dimensions, either by capturing cellular RNAs on barcoded arrays or beads on a surface or delivering barcodes into samples using microfluidics, these methods provide only 2D spatial omics information. The spatial resolution of these methods has been limited by the barcoding grid size (from sub-μm to tens of μm) and diffusion of molecular contents and barcodes (typical to a range that is comparable to the sample thickness, ∼10 μm), making single-cell analysis challenging. The recently developed Slide-tag offers single-cell resolution but still lacks 3D information [5].

Light-controlled reactions offer an alternative approach to spatially dependent barcoding. This new approach of light-directed spatial omics offers a bridge between sequencing-based and imaging-based methods. Because the spatial resolution of this approach is controlled by patterned light, it has the potential for 3D spatial omics with single-cell resolution. For example, the recently reported Light-seq method [6]. demonstrated light-directed, spatially selective barcoding via a photo-crosslinking chemistry that allows DNA barcodes to be linked to nucleic-acid contents in cells or tissues. Another method, ZipSeq [7], tags surface proteins with an antibody conjugated to a photocaged oligonucleotide sequence and introduces spatial barcodes through photo-uncaging and hybridizing a barcode to the uncaged portion of the oligonucleotide. To date, these light-controlled barcoding approaches have been used to provide RNA or protein expression profiling of a small number of spatial regions in the sample. It remains to be demonstrated whether the photochemical reactions used in these methods allow a high degree of spatial multiplexing (namely omics measurements of a large number of spatial regions per sample).

Here, we report MOLseq (Multiplexed optically-controlled ligation followed by Sequencing), a light-directed spatial omics technique. This method uses an optically controlled ligation chemistry to iteratively add short oligonucleotide sequences at specific locations in samples as spatial barcodes, followed by extraction and sequencing of barcoded molecular contents of samples for genome-wide profiling. MOLseq constructs barcodes in a combinatorial manner over multiple rounds, thus providing a highly scalable approach for barcoding such that the barcode diversity grows exponentially with the number of ligation rounds. Barcodes capable of error detection and correction could be used to further increase measurement accuracy. The spatial positions of barcodes are determined by light and, hence, potentially offers a spatial resolution down to a single cell or subcellular level in 3D. We demonstrate that MOLseq can be used to generate a diverse set of barcodes in situ, control the spatial positions of barcodes down to the single-cell level, and provide spatially resolved transcriptomic profiling of cells.

## Results

### Concept of MOLseq for spatial omics

As the first step in MOLseq, we target specific groups of molecules in a sample using a DNA primer, the sequence of which can target, for example, the polyA tail of mRNA, other cellular RNAs, or an oligo-conjugated antibody for proteins. Specifically, in this work, we target the polyA tails of mRNAs by hybridizing them with a poly-T primer, followed by reverse transcription in situ to create cDNAs. We then attach unique oligonucleotide sequences (barcodes) to the cDNA products at different locations in the sample. The spatially barcoded cDNAs are then extracted from the sample and sequenced to determine both the genetic identities and spatial locations of the mRNAs (Figure 1A). As a way to multiplex barcoding, we consecutively attach multiple oligonucleotide sequences to cDNAs at each location in a light-directed manner, each oligonucleotide sequence corresponding to a letter in the barcode. The order of the letters then determines the barcode corresponding to each location.

**Figure 1.**
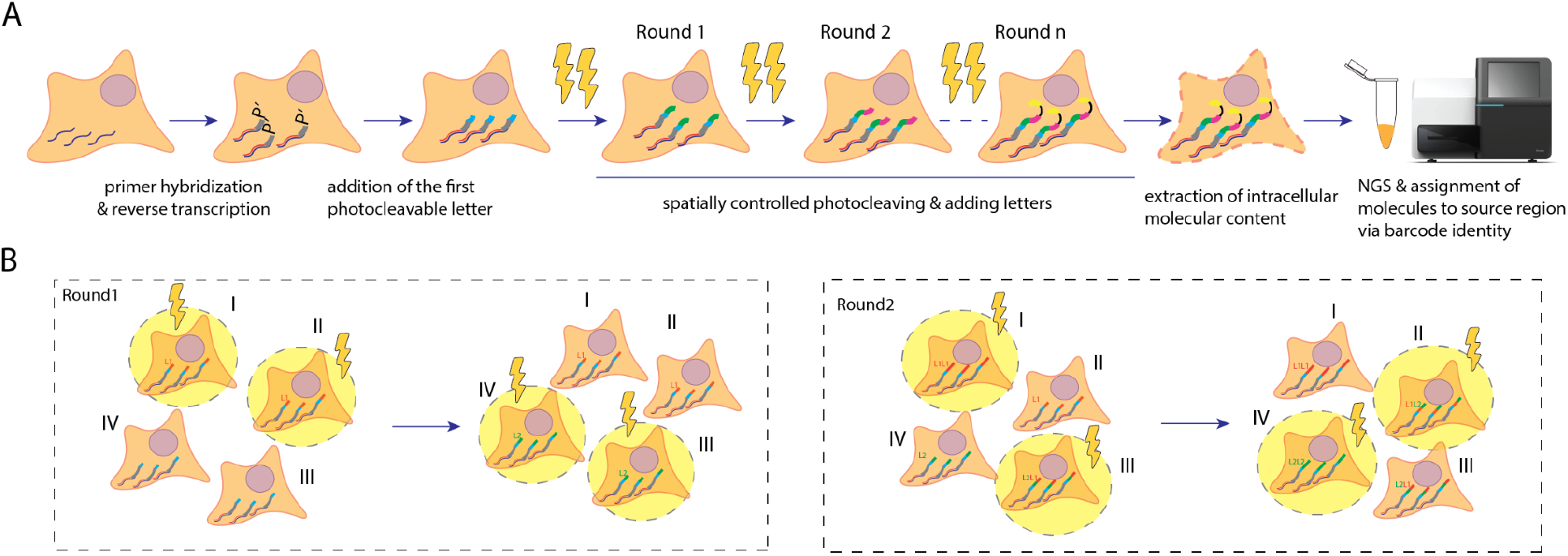
Multiplexed optically controlled barcoding for spatial omics. **A**. MOLseq experimental workflow illustrated in a cell. MOLseq encodes the spatial position of intracellular molecules in the form of oligonucleotide-based barcodes. First, a DNA primer is hybridized to the molecules of interest, such as a poly-T primer targeting the mRNA molecules, followed by in-situ reverse transcription to generate cDNAs. We then build a barcode sequence at the 5’ end of the cDNAs in a spatially dependent manner using light control. Cells are then lysed, and extracted material is sequenced. The barcode sequences allow the assignment of identified molecules to their spatial position, creating a spatially resolved omics map. **B**. Multiplexed generation of spatial barcodes using patterned light. Barcodes are generated by the iterative addition of unique oligonucleotide sequences to the primer in regions illuminated by light. Here, we depict two rounds. Each round completes the addition of one letter to the barcode, either L_1_ or L_2_. Regions I and II, which have letter L_1_ in the first digit position, are illuminated at the same time, followed by the illumination of regions III and IV and addition of letter L_2_ in their first digit. This completes the first round, and we add the second letter to the barcodes in a similar manner in the second round. In this manner, the barcode diversity grows exponentially with the number of rounds, conferring scalability and multiplexity to the design.

Figure 1B illustrates a specific example of such a barcoding process. Suppose we have two letters, L_1_ and L_2,_ and aim to barcode four regions: region I = L_1_L_1_, region II = L_1_L_2_, region III = L_2_L_1_, and region IV = L_2_L_2_. In the first round, regions I and II are illuminated with light and the letter L_1_ is added. Then, regions III and IV are illuminated with light and the letter L_2_ is added. In the second round, we illuminate regions I and III and add an additional letter L_1_ to these regions, followed by illuminating regions II and IV and adding an additional letter L_2_ to these regions to finish the barcoding process. This simple example illustrates the multiplexity and scalability of this barcoding approach: with m letters and n rounds, we can generate *m*^*n*^ unique barcodes in *m* x *n* letter-ligation steps (i.e., *m* steps for each of the *n* rounds; spatial locations encoded with the same letter in the same round can be illuminated simultaneously). Moreover, the barcoded regions do not have to cover the whole sample and can be chosen based on interest.

At the end of the barcoding process, each cDNA contains both the spatial barcode and transcript identities. We then extract cDNAs from cells and processed them for next-generation sequencing (NGS) (Figure 1A). A variety of NGS platforms are available, which offer different sequencing depth and read length options. An optional 5’ enrichment step can be implemented during library preparation for platforms with short read length or limited depth.

### Light-controlled in-situ barcode generation

We generate light-controlled barcodes through recursive ligations of letter-representing DNA sequences with a photocleavable spacer (PC) linked to another oligonucleotide sequence (Figure 2A) [8, 9]. The process begins with a primer that has a 5’ phosphate group, to which we ligate a letter using a ligase enzyme. A splint is chosen to facilitate the ligation, with a complementary sequence to a part of the current barcode sequence and a part of the incoming letter. The ligation product is then exposed to UV light to photocleave the PC and expose a 5’ phosphate group, allowing the next round of ligation to occur, as illustrated in Figure 2A. If the letter is not photocleaved, there is no 5’ phosphate group, preventing further ligation. In a proof-of-principle experiment with two rounds of ligations, in which one of the samples was exposed to UV after the first round of ligation of letter L_1_, and the other sample was not, we indeed observed ligation of letter L_2_ only to the sample that was exposed to the UV light, but not to the other sample (Figure 2B).

**Figure 2.**
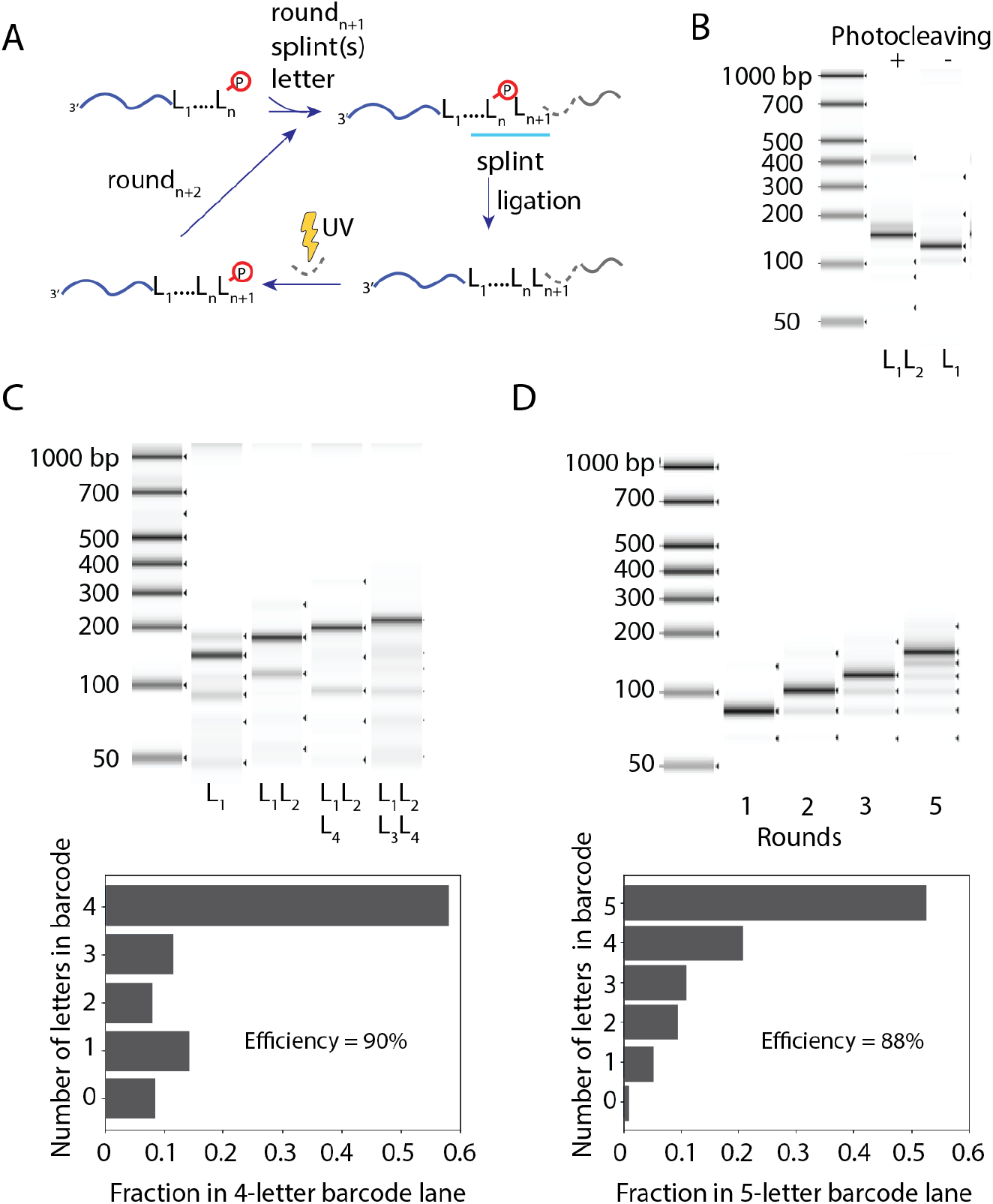
Iterative light-controlled ligations of letter-representing DNA sequences with a photocleavable spacer. **A**. A barcode is constructed from a sequence of letters L_1_, L_2_,… Suppose that *n* letters have already been added with a phosphate group at the 5’ end. The next letter (gray) and a splint (blue) are added to enable the ligation reaction. A splint is a 15-base sequence, five and ten of which are complementary to the incoming letter and the most recently ligated letter, respectively. The incoming letter has a photocleavable spacer (PC, shown in dashed lines) followed by another oligo sequence without a 5’ phosphate group. If this newly extended barcode is exposed to UV light, the PC group will be cleaved, exposing a free 5’ phosphate group, which will allow for the next round of ligation. **B**. Gel image of the extracted material from two samples, one of which has been exposed to UV light after the first round of ligation (middle lane labeled with L_1_L_2_) and the other one has not (right lane labeled with L_1_), preventing the addition of the second letter and demonstrating optical control of this ligation process. **C**. Top, sequential ligations showing the construction of a 1, 2, 3, and 4-letter barcode (lanes labeled with L_1_, L_1_L_2_, L_1_L_2_L_4_, and L_1_L_2_L_3_L_4_, respectively). Bottom, Gel quantification of the 4-letter barcode construction case (right-most lane of the above gel). We estimate the efficiency of each round of letter ligation by taking the fraction of the correct length barcode to the power of 1/*n*, where *n* is the number of rounds (*n* = 4 in this case). Shorter species of 0,1,2 or 3-letter barcodes are due to incomplete photocleavage and ligation. The calculated efficiency per round here is 90%. **D**. Top, to demonstrate barcode multiplexity, we use all four letters with all the necessary splints for each round of letter addition in the construction of a 1, 2, 3, and 5-letter barcode (lanes labeled with 1, 2, 3, and 5, respectively). Bottom, Gel quantification of the 5-letter barcode construction case (right-most lane of the above gel). The average efficiency per round is estimated to be 88%. Left lane in (B, C, and D) is the molecular weight standard.

Next, we demonstrated that this process can be used to continuously add multiple letters with high efficiency by generating 1, 2, 3, and 4-letter long barcodes (each letter 20 bases long), in cultured Human Osteosarcoma (U2OS) cells (Figure 2C). In this case, to facilitate quantification of the barcoding products by gel electrophoresis, the poly-T primer was not reverse transcribed to preserve the original sequence length and had a PCR primer at the 3’ end to allow PCR amplification. After the light-controlled ligation steps, the barcoded primer was extracted, and a PCR was performed for one cycle to generate a second strand. Then, the strand was analyzed by gel electrophoresis. From the ratio of the intensity of the correct size band to smaller bands arising from inefficient photocleaving or ligation, we estimated the efficiency of each round of letter addition in the 4-letter long barcode to be 90%.

When generating distinct barcodes in parallel, we need to be able to add a letter to any possible combination of existing letters on the cDNAs to ensure that this process is multiplexable. To achieve this, we could use a pool of splints, each of which has complementary sequences to the incoming letter and one of the previously added letters. Figure 2D demonstrates barcode generation by ligating four different letters with all possible splints over 1, 2, 3, and 5 rounds with a pool of splints that could facilitate ligation of all 4 incoming letters and all prior letters (namely a total of 4 x 4 splint sequences). Similar to the estimation of efficiency in the single barcode case described above, we estimated the efficiency of letter addition per round to be 88% for this 5-letter barcoding process by comparing the intensity of the correct size band to smaller bands on the gel. We then sequenced the 5-letter-long barcode and detected 795 unique barcodes. This number is less than the expected number (4^5^ = 1024) of total possible barcodes. Since this experiment had all the 4 letters present in every round, and different letters might compete with each other, resulting in a reduced number of barcodes generated. We anticipate the barcode diversity to be higher in the actual MOLseq barcoding process, where only one incoming letter would be present for ligation at any time without potential competition from other letters.

### Light-controlled barcoding in a spatially resolved manner

We generate spatially selective barcodes by directing patterned UV light onto the sample using a Digital Micromirror Device (DMD). Using poly-T primers attached to mRNAs in cultured U2OS cells as a model system and fluorescently labeled FISH probes that target ligated letters, we demonstrated the ability to achieve spatially controlled photocleaving and ligation on selected sample areas ranging from hundreds of microns down to a single cell (Figure 3A, B).

**Figure 3.**
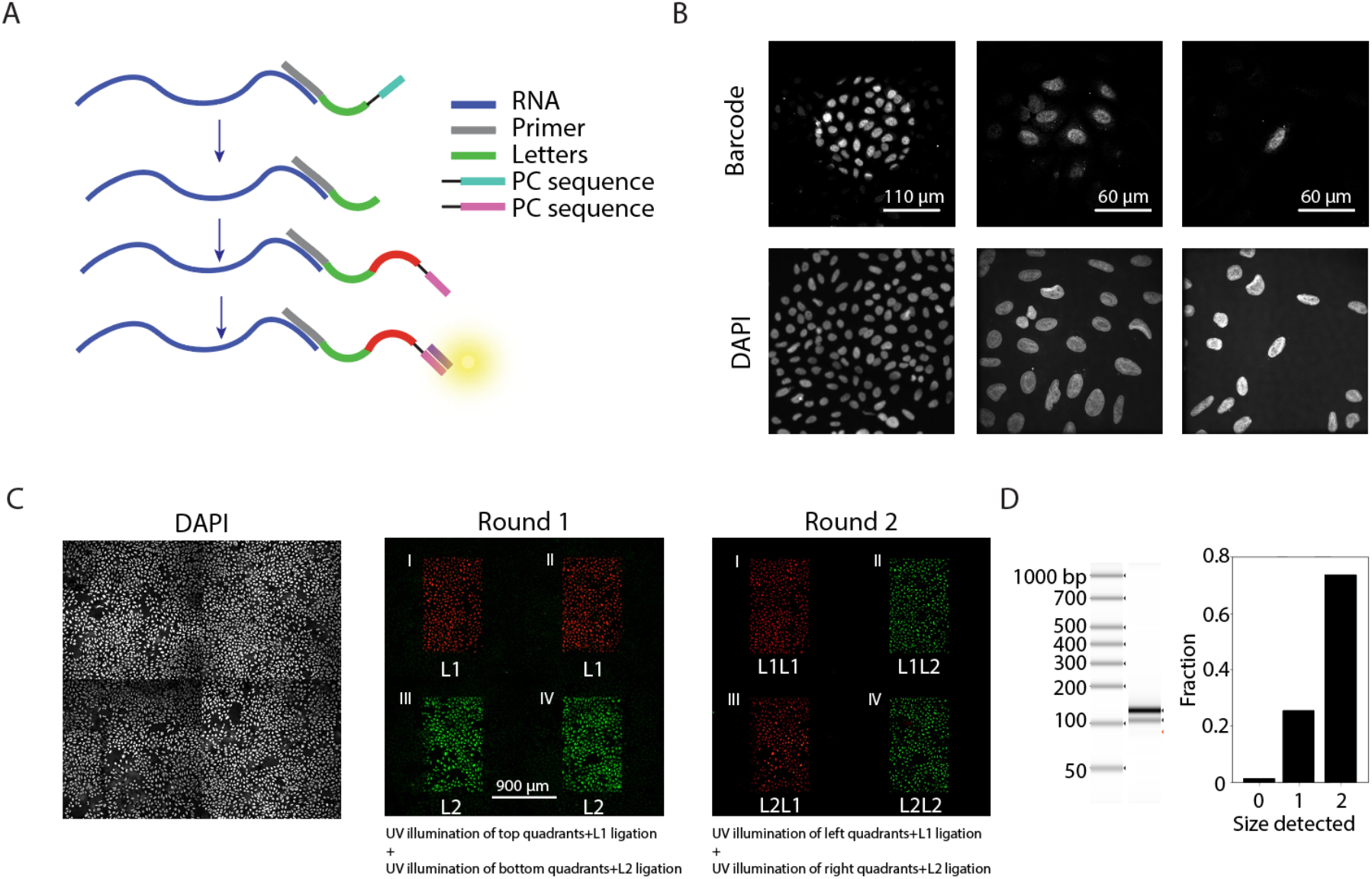
Spatially controlled in-situ barcode generation via patterned light using DMD. **A**. Detection scheme of the individual letters in the barcode sequence. A poly-T primer (gray) is hybridized to the mRNA molecules (blue), and a letter with a PC (green and cyan) is ligated to the primer. DMD is used to photocleave the PC in an optically defined region, generating a 5’ phosphate for further ligation, and a ligation mixture with the next letter (red and pink) is added. Successful ligation of the letter to the barcode is detected through hybridizing a FISH probe complementary to the sequence next to the PC. **B**. Spatially controlled letter addition to specific regions in the sample of 220 µm, 90 µm, and 20 µm in diameter. FISH signal coming from spatially controlled ligation product in the 220 µm, 90 µm, and 20 µm diameter circles and the 4’,6-diamidino-2-phenylindole (DAPI) signal coming from all cells in the field of view are shown on the top and bottom panels, respectively. **C**. Construction of four unique barcodes in four different sample regions over two rounds of ligation. The barcode is constructed from two letters, L_1_ and L_2_, detected with Alexa647 (red) and ATTO565 (green) FISH probes, respectively. In the first round, the two regions on the top (regions I and II) are illuminated first to enable ligation of the letter L_1_. Afterwards, the bottom two regions (regions III and IV) are illuminated to enable ligation of L_2_. In the next round, the same steps are repeated with a different illumination pattern where the two ligation reactions are preceded by the illumination of the regions I and III first and regions II and IV. This experimental scheme generates four unique barcodes L_1_L_1_, L_1_L_2_, L_2_L_1_, and L_2_L_2_ in regions I, II, III and IV, respectively. DAPI image of all cells are shown in the left panel. **D**. Left: A gel image of the extracted material from the sample described in (C). Left lane: molecular weight standard. Right lane: extracted sample material. Right: Quantification of the relative intensities of each band in the sample-material lane, demonstrating that 75% of the primers are extended by the complete barcode length consisting of two letters.

This process of spatial barcode generation can be repeated over multiple rounds to build diverse barcodes in various regions. We demonstrated the creation of four distinct barcodes in four distinct spatial locations using two letters, L_1_ and L_2_, which can be detected using two differently colored fluorescent probes (false colored red and green for L_1_ and L_2_, respectively, in Figure 3C). In the first round, we added the letter L_1_ to two sample regions (regions I and II) and the letter L_2_ to two other regions (regions III and IV). In the next round, we again ligated L_1_ and L_2_, but to different region combinations (L_1_ to regions I and III and L_2_ to regions II and IV). After each round, we detected the ligation product by FISH probes that were complementary to the sequence 5’ end of the PC moiety in the letter-representing oligonucleotide (Figure 3A). Indeed, FISH imaging verified the successful ligation of letters L_1_ and L_2_ in the designed order in four sample regions in a multiplexed manner (Figure 3C).

As a further validation, we ligated a PCR primer onto the four regions, extracted the barcoded primers, PCR-amplified them, and analyzed them by gel electrophoresis. We primarily observed products with a 2-letter long barcode (75%) and only a small fraction with a 1-letter barcode (25%) (Figure 3D). The presence of 1-letter barcode products may arise from less than 100% photocleavage efficiency in the selected region or photocleaving and ligation products outside the region of interest. The efficiency here is lower than the experiments described in Figure 2, for which we used a Light Emitting Diode (LED) to illuminate the sample and perform photocleaving uniformly. In contrast, here, we used DMD to direct light from a different LED source to the sample in a spatially controlled manner. The light intensity at the sample in the latter case was lower, and with the optimization of light intensity, we anticipate that high letter addition efficiency with light directed by DMD will be also possible, as in the case of using uniform illumination.

### Spatially resolved transcriptomics by light-controlled barcoding

As a proof-of-principle validation of the ability of our light-directed barcoding approach to provide spatially dependent transcriptomic profiling, we performed a mixed-species experiment using U2OS and Mouse Embryonic Fibroblast (MEF) cell lines, plated in two separate regions of the same coverslip. This approach allowed us to create distinct but adjacent regions of cells of each species without mixing. We employed the full MOLseq process to barcode and sequence this sample (Figure 4A). We first hybridized poly-T primers to the sample, performed reverse transcription to extend the primers, and then barcoded cDNA products in the two different regions each with a unique 2-letter barcode, namely L_2_L_4_ for the U2OS cells on the left side and L_1_L_3_ for the MEF cells on the right side of the sample. Building barcodes with more than one letter that are distinct between the two regions introduces error-detection capability and increases the accuracy of spatial barcoding. Approximately 3,000 cells were barcoded in each species. We confirmed the generation of 2-letter barcodes in a region-dependent manner by detecting the presence of the expected barcodes using FISH with differently colored probes for different barcodes (Figure 4B). We found that each barcode’s fluorescent signal was highly specific to its targeted region, where the off-target barcoding rate was only 4% for L_2_L_4_ and 1% for L_1_L_3_ (Figure 4B).

**Figure 4.**
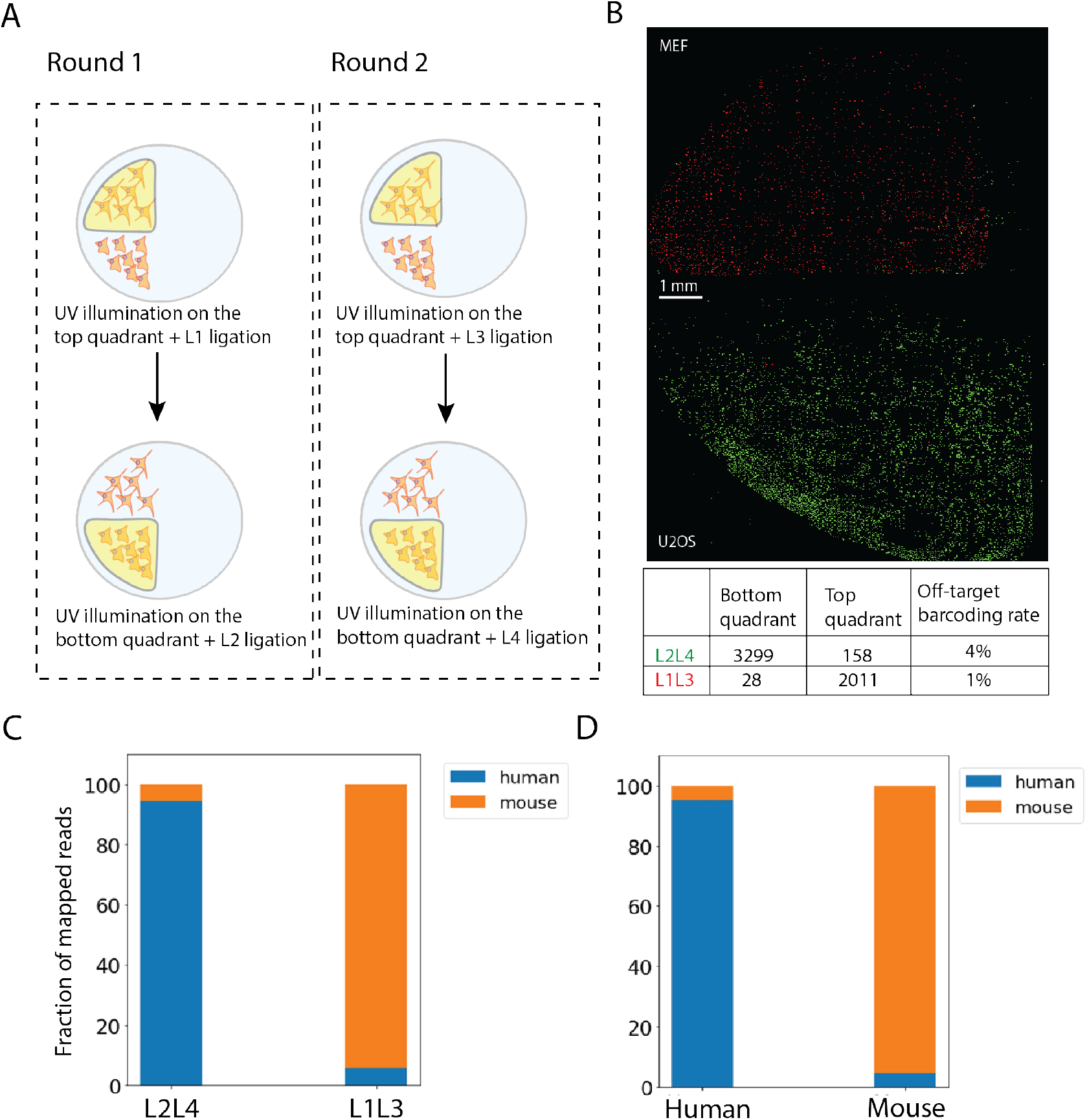
Spatially resolved transcriptomics by light-controlled barcoding with MOLseq. **A**. Mouse and human cell lines are plated on the top and bottom sides of the sample, respectively, and four letters are used to construct two unique barcodes encoding the two regions (L_1_L_3_ for the top region) and (L_2_L_4_ for the bottom region). **B**. Image of the U2OS and MEF cells hybridized with FISH probes complementary to half of each letter in the barcode sequence. FISH probes for the barcode L_2_L_4_ is labeled with Alexa647 (green) and FISH probes for L_1_L_3_ is labeled with ATTO565 (red).The table at the bottom shows the quantification of number of green and red cells in each quadrant. The off-target barcoding rate is calculated as the ratio of the number of cells with red or green fluorescence on the untargeted quadrant to all of the red or green cells. **C**. Fraction of the sequencing reads containing the specified spatial barcodes that are mapped to the human and mouse genome. For the barcode L_2_L_4_, 95% of all mapped reads are human, and for the barcode L_1_L_3_, 96% of all mapped reads are mouse. **D**. Species distribution of reads generated from U2OS and MEF cells cultured in physically separate chambers and processed independently until the sequencing stage. We still see a ∼5% cross-species mapping.

We then extracted the cDNA content, enriched for the 5’ end containing the barcode and part of the transcript sequence, and performed short read sequencing with paired-end reads, generating a total of ∼600K reads. The reads were mapped to the mouse and human genomes, and the intronic and intergenic reads were filtered out. For the barcode L_2_L_4_, the relative fractions of reads mapped to human and mouse species were 95% and 5%. For the barcode L_1_L_3_, the relative fractions mapped to human and mouse species were 4% and 96% (Figure 4C). When we cultured U2OS and MEF cells in physically separate chambers and processed them separately until the sequencing stage, we still observed a ∼5% cross-species mapping rate (Figure 4D), presumably due to a certain level of homology that exist between mouse and human transcriptomes, consistent with results from previous studies [10]. Hence, the error rate in MOLseq for spatially resolved transcriptomics is likely substantially less than 5%.

## Discussion

In this work, we developed MOLseq as an approach that brings together the power of optical control and sequencing for spatial omics in a multiplexed and scalable manner. The iterative in-situ barcode construction, using light-directed cleavage and ligation of oligonucleotides, allows for an exponential increase in the diversity of barcodes, which should allow high-throughput spatial omics measurements across a wide range of length scales, from single cells to tissue regions. The flexibility and versatility in light patterning should allow the spatial resolution and number of targeted spatial regions to be readily adjusted based on experimental needs. Using light patterning in 3D, this method could also be extended to allow genome-wide, 3D spatial omics.

We demonstrated in cultured cells that we can perform in situ light-controlled DNA cleavage and ligations with up to 90% efficiency. Using a pooled mixture of splint sequences to allow any letter-representing oligonucleotide sequence to be added to desired locations independent of the previously added letters, we can construct a high diversity of barcodes in a time-efficient manner. Indeed, we generated created ∼800 unique barcodes with only 5 rounds of ligations using pooled mixtures of barcode letters and ligation splints, demonstrating that this multiplexed, light-controlled ligation is a viable approach for scalable spatial barcode generation. The multi-letter-based barcode design not only allows for scalability but also provides error detection and correction capability. Using a DMD to select photocleaving and ligation regions at various length scales, we demonstrated optically controlled barcoding in regions ranging from millimeters to a single cell and obtained high spatial specificity. Combining this in-situ optically controlled barcoding with sequencing, we demonstrated spatially resolved transcriptomic profiling of cells with high accuracy. The data presented here focuses on mRNA measurements, but MOLseq barcoding can be adapted to measure other molecular species, such as detecting protein molecules by oligonucleotide-conjugated antibodies [11-14].

MOLseq also has its limitations. Since it relies on photocleaving by patterned light and subsequent ligation for spatial selectivity, off-target photocleaving from the tails of patterned UV light or background cleavage can introduce errors, namely crosstalk of barcodes between different spatial regions. This could be especially prominent when addressing small spatial regions. In this work, we performed high-accuracy spatial transcriptomics measurements on relatively large sample regions and have yet to find out what the error rate is when measuring spatial regions as small as a single cell. Moreover, the illumination crosstalk can cause correlated errors, which can be difficult to mitigate with error correction. In addition, the barcode diversity generated by MOLseq is limited by the photocleavage and ligation efficiency. With the current 90% efficiency per ligation round, it would be reasonable to limit the number of ligation rounds (*n*) to ∼5. In order to generate a diversity of 10^5^ barcodes (i.e., measure 100,000 spatial regions simultaneously), we will need the number of distinct letters (*m*) to be 10 because the barcode diversity is equal to *m*^*n*^. Since we construct the barcodes by ligating one letter at a time, this would require 10 ligation steps per round, and hence 5 × 10 = 50 total ligation steps to generate the barcodes, which will take ∼3 days based on our current protocol. It is worth noting that an increase in *n* would rapidly increase the barcode diversity with a minimal effect on the total barcode generation time if we keep *m* × *n* constant by reducing *m* accordingly (for example, 5^10^ is approximately 10 million). We anticipate that future work to improve the ligation efficiency will allow *n* to be increased, hence increasing the time efficiency and diversity in barcode generation, which may ultimately allow comprehensive, genome-wide spatial omics analysis of large samples at single-cell resolution.

## METHODS

### Sample preparation

Human Osteosarcoma (U2OS) or Mouse Embryo Fibroblast (MEF) were cultured and plated onto Ibidi µ-Slide VI 0.4 lanes for barcoding experiments without reverse transcription and sequencing and Ibidi Culture-Insert 4 Well in µ-Dish 35 mm for the co-culture and MOLseq experiment. The Ibidi chambers were incubated with Poly-D-Lysine solution (0.1 mg/ml, Thermo Fisher, A3890401) for one hour and washed with water before use. The cells were plated and incubated at humidity-controlled incubators at 37 °C overnight to allow proper attachment and flattening and fixed with 4% paraformaldehyde (PFA) diluted in 1x Dulbecco’s phosphate-buffered saline (DPBS). We then hybridized the cells with the poly-T primer (**Table S1**) at 1 uM concentration in 2x SSC at 37 °C overnight and washed the unbound and non-specifically bound primers with 30% formamide in 2x SSC at 47 °C for 30 min twice followed by 1x DPBS twice, Ambion nuclease-free water (Thermo Fisher, AM9920) twice, and 2x SSC twice. The poly-T primer was replaced with a poly-T primer containing locked thymidine bases for experiments in Figure 2. The samples for which reverse transcription was needed were incubated with reverse transcription (RT) buffer composed of 1.875 mM dNTP mix (25 mM each) (Thermo Fisher, R1121), 1x Maxima RT buffer (Thermo Fisher, EP0742), 600 units/ml RNAse inhibitor, murine (NEB, M0314S), 600 units/ml RNasin® Plus RNase Inhibitor (Promega, N2611), 2.5 µM Template Switching Oligo (TSO) (**Table S1**), 10 units/µl Maxima H Minus Reverse Transcriptase (Thermo Fisher, EP0751) at 37 °C overnight and washed with 2x SSC twice and 1x PBS twice to remove excess reagents.

### Photocleaving and ligation of letters for barcode generation

We prepared 2x ligation stock buffer weekly, which was composed of 132 nM Tris-HCl, 20 mM MgCl_2_, 15% Polyethylene glycol (PEG 8000) at pH 7.6 and stored at room temperature. Immediately before use, we supplemented the stock buffer with 20 mM ATP (Thermo Fisher, R0441) and 20 mM DTT (NEB, 7016L), both of which have only been freeze-thawed once. Samples were washed with 1x ligation buffer twice and incubated with the ligation mixture containing 1x ligation buffer, 0.1 % Triton-X 100 (Sigma-Aldrich, 93443), 0.4 µM splint(s), 0.4 µM letter-representing DNA and 40 U/µl Quick ligase enzyme (NEB, M2200S) for 1 hour at room temperature in dark. We selected letter sequences to have at least 40% GC content with no G or C at the ends of the sequence to support high ligation efficiency [15]. The splints contained sequences complementary to the five bases at the 3’ end of the incoming letter and the ten bases at the 5’ end of the prior letter. The splints are named by using a numbering convention of incoming letter ID followed by the prior letter ID on the barcode. For example, ligation of L_1_ to L_2_ would be facilitated by the splint12. All letter and splint sequences are shown in **Table S1**.

Following the ligation, samples were washed with 1x PBS twice, 30% formamide in 2x SSC twice, and nuclease-free water twice to remove excess reagents. Photocleaving for the ligation product was performed using a stand-alone LED light source, 365 nm, or an inverted Olympus IX-71 microscope with a 10x objective coupled to Mightex 1000 series polygon1000G model DMD with a different LED source, 365 nm. We illuminated the desired regions of each sample at an intensity of 3 mW/mm^2^ for 30 seconds with the DMD or 10 mW/mm^2^ for 30 seconds with the LED and washed the cleaved products with 1x PBS twice and nuclease-free water twice.

### FISH detection of light-controlled barcode letter additions

U2OS cells were prepared as described in the “Sample preparation” section. After fixation, we ligated the entire sample with the PC-containing letter L2 before starting our barcoding ligations such that the whole sample is photocleavable. We then washed the sample in wash buffer composed of 30% formamide in 2x SSC twice and 1x PBS twice. We then incubated the cells with phosphatase buffer composed of 1x rCutsmart buffer (NEB, B6004S), 0.1 % TX-100, 250 U/ml quick CIP enzyme (NEB, M0525S) for 20 min at 37 °C to remove any free 5’ phosphate groups that may have formed due to stray light and washed the samples with 1x PBS once and nuclease-free water once. This ensures that only regions that are photocleaved by light participate in subsequent ligation steps. We illuminated the regions (10% of total cells) of interest using patterned light generated by DMD and proceeded with the ligation of letters. To allow detection of the barcodes by FISH in this experiment, we used letters contained the TGCGAACTGTCCGGCTTTCA or GATCCGATTGGAACCGTCCC next to the PC. This sequence was then detected with a complementary probe conjugated to Atto 565 (FISH probe 1) or Alexa 647 (FISH probe 2) with a thiol linkage. After the ligation of L1 and L2, we incubated the samples with the FISH probes at a 10 nM final concentration in wash buffer. We removed the excess reagents by incubating the samples in wash buffer for 10 min once and nuclease-free water once. The samples were imaged with an inverted Olympus IX-71 microscope using 560 nm and 647 nm excitation light. We observed that the signal coming from the FISH probes did not go away after photocleaving and extensive washing. This is unlikely to be due to inefficient photocleaving, as that would prevent further ligation, but we observed that the vast majority of the ligation products after the first round were capable of ligation in the second round. It is possible that after photocleaving, the FISH probed labeled photocleavage product remained stuck on the sample. To overcome this problem, we used FISH probes conjugated to fluorophores by a disulfide bond. Following the detection of the first letter, samples were subjected to tris(2-carboxyethyl)phosphine (TCEP) incubation, cleaving the disulfide bond and removing the fluorophores before moving forward. We incubated the samples in wash buffer supplemented with 0.5 M TCEP and blocker sequences at 10 nM (blocker 1 and blocker 2, **Table S1**) for 20 min at room temperature. Following this, the samples were washed with 30% formamide in 2x SSC for 10 minutes and then twice with 2x SSC. Letters ligated in the next round were detected the same way and then treated with phosphatase, as described earlier. We then photocleaved the PC and performed a final round of ligation of the PCR primer.

### FISH detection of light-controlled barcodes in the co-culture experiment

In these experiments, we perform FISH labeling in two steps, the first step with an adaptor probe that contains both a target sequence complementary to a barcode letter and a readout sequence that can be detected by a complementary, fluorescently labeled readout probe. We used adaptor probes complementary to part of L_1_L_3_ (Barcode 1 adaptor probe sequence, Table S1) and L_2_L_4_ (Barcode 2 adaptor probe sequence, Table S1) flanked by readout sequences to detect the barcodes in the sample. After barcoding was completed, we incubated the samples with the adaptor probes at 0.1 µM concentration diluted in wash buffer composed of 30% formamide, 2x SSC, for 20 min at room temperature followed by a 20 min wash step with the wash buffer to remove the non-specifically bound and excess reagents. We then incubated the samples with the readout probes at 10 nM final concentration diluted in wash buffer for 20 minutes and washed the excess reagents with the wash buffer for 10 minutes. All incubation and wash steps were performed in the dark. For quantification, images are thresholded to remove background and analyzed using the “Analyze particles” module in ImageJ to determine the number of fluorescently labeled cells.

### Extraction and detection of barcoded primer or cDNA for barcode-size measurements

To extract the primer or cDNA from the samples, we heated the samples in 50 ul water supplemented with 0.1 % Triton-X 100 to 93 °C for 4 min and immediately aspirated the solution into an Eppendorf tube and left it at room temperature for several minutes for it to cool down. For Figures 2B and 2C, we generated a second strand by performing one PCR cycle using only the reverse primer with Seqamp DNA polymerase (Takara Bio., 638504) in SeqAmp CB PCR buffer (Takara Bio., 638526) according to the manufacturer’s instructions. For Figures 2D and 3D, we performed 5-10 rounds of qPCR using both forward and reverse primers. We then purified the PCR product using AMPure XP Bead Based Reagent (Beckman Coulter, A63880) at 0.8x v/v ratio according to the manufacturer’s instructions. The purified product was run in a High Sensitivity D1000 ScreenTape (Agilent, 5067-5584) using the Agilent 2200 TapeStation system.

### Library preparation and RNA sequencing analysis

For samples used for sequencing, we ligated SmartSeq3 sequences to the barcoded cDNA. We extracted the cDNA similarly as described in the previous section and amplified the extracted material using long-distance (LD) PCR with Seqamp DNA polymerase according to the instructions outlined in Takara Smartseq V4 kit using a forward sequence complementary to Smart seq3 and a reverse primer complementary to the TSO sequence. The amplified product was purified using Ampure XP beads at 0.8x v/v ratio. We used the Nextera XT DNA Library Preparation Kit (Illumina, FC-131-1024) for tagmentation with a few modifications. First, we amplified the tagmented products with our custom forward and reverse primers (Table S1) for 5’ enrichment. Following the purification of the PCR product using Ampure XP beads at 0.8x v/v ratio, we performed another PCR round to add the Illumina P5, i5, P7, and i7 sequences for downstream sequencing and purified the products using Ampure XP beads at 0.8x v/v ratio. Sequencing was performed in the Miseq system using the MiSeq Reagent Kit v3 (150-cycle) (Illumina, MS-102-3001). Following sequencing and demultiplexing, the reads were filtered for having the right barcode sequence and then mapped onto the appropriate genome using STAR [16]. The aligned reads from STAR were enumerated using RSEM [17].

## Note

We note that a similar approach is independently developed and reported in bioRxiv recently: Battistoni et al., https://www.biorxiv.org/content/10.1101/2024.05.20.595040v1.

## Acknowledgements

We thank members of the Zhuang lab for helpful discussions. This work is in part supported by the National Institutes of Health. X.Z. is a Howard Hughes Medical Institute Investigator.

## Supplementary Materials

**Table S1.**
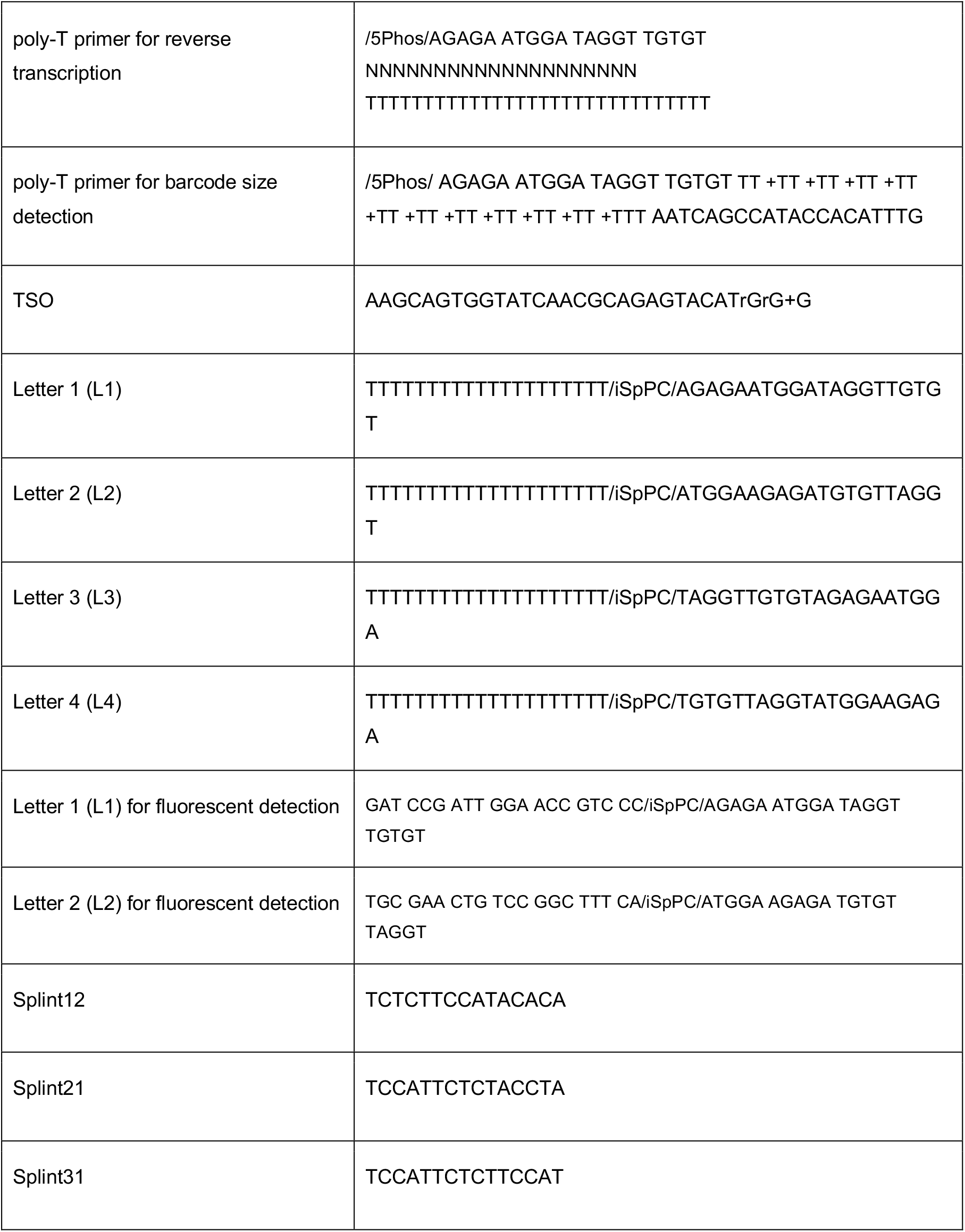
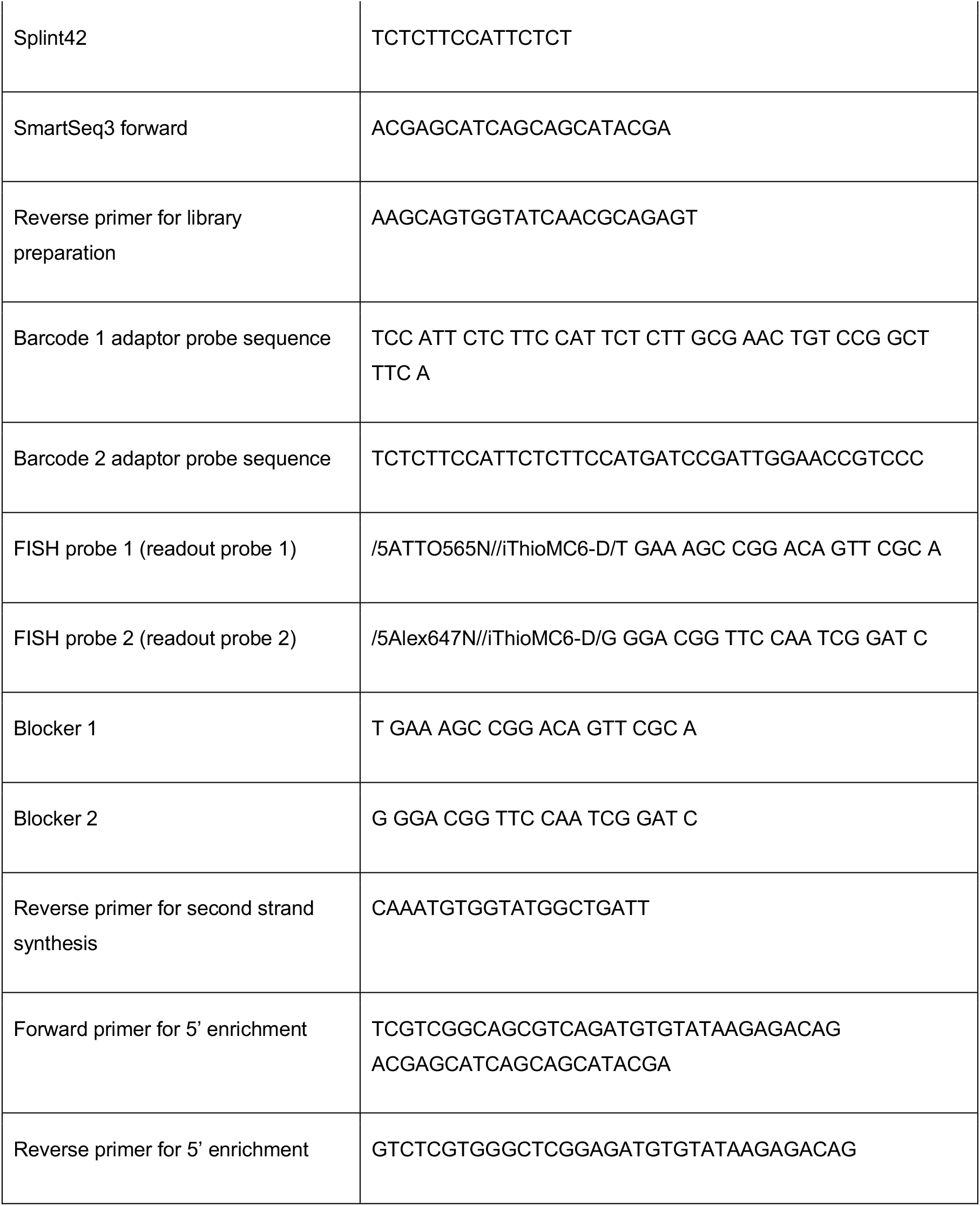
Sequences of primers, letters, and splints used in this study. All oligonucleotides are purchased in lyophilized form from Integrated DNA Technologies (IDT).

